# A phage-derived reconfigurable effector associated with an actinobacterial contractile nanomachine tailors bacterial responses to competition

**DOI:** 10.1101/2025.11.06.686954

**Authors:** Toshiki Nagakubo, Tatsuya Nishiyama, Shumpei Asamizu, Hiroyasu Onaka, Nobuhiko Nomura, Masanori Toyofuku

**Affiliations:** Institute of Life and Environmental Sciences, University of Tsukuba, Tsukuba, Japan; Tsukuba Institute for Advanced Research (TIAR), University of Tsukuba, Tsukuba, Japan; Microbiology Research Center for Sustainability (MiCS), University of Tsukuba, Tsukuba, Japan; Life Science Research Center, Nihon University, Fujisawa, Japan; Engineering Biology Research Center, Kobe University, Kobe, Japan; Graduate School of Agricultural and Life Sciences, The University of Tokyo, Bunkyo, Japan; Faculty of Science, Gakushuin University, Toshima, Japan; Life Science Center for Survival Dynamics, University of Tsukuba, Tsukuba, Japan

## Abstract

Contractile injection systems (CISs) are derivatives of phage tails and widely distributed in prokaryotes. CISs load cognate effectors and eject them through contractile actions resembling those of phage tails. Ejected effectors play central roles in CIS functionality by acting on target cells and mediating various biological processes. Here, we report a novel group of CIS effectors related to phage tapemeasure protein, the transmembrane component of the phage infection machinery. This group is broadly distributed within the class actinobacteria, one of the bacterial classes in which CIS gene clusters are highly conserved, and is represented by Sle1, a cognate effector of the intracellularly localised *Streptomyces lividans* phage tail-like nanoparticle (SLP). This effector is associated with Sle2, which contains a CIS effector core domain and interacts with the SLP core component. Sle1 is packaged inside SLP and is translocated to lipid membranes along with SLPs. The functional domain of Sle1, probably through interactions with ribosome-containing subcellular fractions, upregulates the membrane-associated proteome in *S. lividans* and *E. coli*. This effect modifies the physiological properties of *S. lividans*, ultimately enhancing its adaptation to microbial competition. In addition, we revealed that Sle1-type effectors conserved among actinobacterial species are structurally and functionally diverse in their functional domains. One of them from *Micromonospora eburnea* constitutes a novel toxin-antitoxin system and introducing its functional domain into Sle1 reprogrammes the phenotypic responsiveness of *S. lividans* to neighbouring bacteria. Our findings illustrate that phage elements can be incorporated into CISs as reconfigurable platforms for bacterial adaptation to various environmental conditions.

**Importance:** Bacterial CISs have attracted interests for their importance in microbial ecology and potential in biotechnological applications. However, understanding of their functional diversity is currently limited because many CIS effectors remain unannotated due to a lack of inferable structural and genetic signatures. Our findings on Sle1 and its relatives illuminate a previously unidentified class of CIS effectors with phage tapemeasure protein-related modular architecture, association with the CIS effector core domain, and wide distribution within the major class of actinobacteria, substantially expanding the known repertoire of effector classes. The impact of Sle1 on *S. lividans* suggest a link between CIS effectors and bacterial adaptation to environmental conditions, highlighting unexplored functional diversity of CIS effectors as tuners of bacterial phenotypes in communities. This work offers routes to manipulate ecological behaviours of actinobacterial species and to access their cryptic traits through effector modulation.

## Introduction

Bacteria and phages, viruses that infect bacterial cells, are involved in a continuous arms race. Although recent studies have characterised various systems that protect bacterial cells from phage infection and its detrimental ecological consequences by eliminating phage elements, bacteria often maintain genes encoding phage tail-like nanostructures within their genomes (1–4). Such nanostructures are no longer infectious and can be ecologically beneficial to the host bacteria under certain circumstances (5,6).

Contractile injection systems (CISs) are a group of prokaryotic phage tail-like nanostructures with macromolecular structures and mechanisms of action resembling those of the contractile tails of myophages. Both CISs and myophages are composed of a central, hollow tube encased by sheath that contracts during action, with a baseplate complex connecting them to spike. On the basis of the conservation of structural characteristics, CISs have been proposed to have evolutionarily diverged from tailed phages (7), perhaps through evolutionary events that caused genetic immobilisation. While phages inject nucleic acids into target cells through tail contraction, CISs are not associated with viral genetic material but load certain effectors inside the tube lumen and eject them through contractile action similar to that of phages. The ejected CIS effectors can act on target cells through their enzymatic activities or other unknown mechanisms (8,9). Because of their sophisticated structures and functional versatility, CISs have been employed by a wide range of prokaryotic species as mediators of various biological processes (8–10). At present, known CISs and CIS-related nanostructures can be grouped into two classes based on their localisation. Some CISs are extracellularly released, after which effectors are injected into target cells by attaching their fibers to the cell surfaces (8–11). Others are intracellularly localised and several studies have implied their potential evolutionary relationship with type VI secretion systems, a class of gram-negative bacteria-specific phage tail-like nanomachines (12). Although the former class of extracellular CISs has been relatively well-investigated and characterised, the latter class of intracellular CISs remains poorly understood, particularly in terms of their cognate effectors. Given the rapidly increasing interest in the medical and biotechnological applications of CISs through the modification of effectors (13,14), a comprehensive understanding of the natural effector repertoire is required.

*Streptomyces* has long been investigated as a producer of various bioactive metabolites, including antibiotics, and is notable for its highly conserved CIS-related gene clusters (15). In our previous studies, we identified and characterised a *Streptomyces lividans* phage tail-like nanoparticle (SLP) that is closely related to CISs and was originally identified in *Streptomyces lividans* TK23 (16,17). SLP represents the most widely conserved class of actinobacterial CIS-related nanostructures, including the CIS*^Sc^* of *S. coelicolor*, which is a close relative of *S. lividans* (18). The SLP gene cluster is located in the genetic region adjacent to threonine-tRNA, implying that these CIS-related nanostructures are derivatives of an ancient phage infecting bacteria using tRNA loci as attachment sites (19). In the *Streptomyces* life cycle, which encompasses vegetative growth of substrate mycelia and subsequent aerial mycelial erection and spore formation, SLPs are mainly produced during the vegetative growth (17). In addition, SLPs show unique intracellular localisation potentially associated with a complex protein-protein interaction network involving ribosomal proteins, and their involvement in microbial interactions has been suggested (16,17). Although these observations suggest a potential role for SLPs in the central cellular systems, the cognate SLP effector and its function remain unclear.

Here, we report the identification of a cognate effector of SLP and its effect on the producer bacterium. We show that the cognate SLP effector represents a novel class of CIS effectors, and that its functional domain broadly affects the cellular functions of *S. lividans* by modulating cellular protein profiles. We also propose that the functionality of these effectors was acquired and diversified from the phage infection machinery to foster the adaptation of producer bacteria to varying environmental conditions.

## Results

### Identification of the SLP cargo

In our previous analysis to identify SLP-associated proteins (17), we noticed that several proteins with unknown functions were identified as being associated with SLPs (Supplemental Fig. 1). Among these, the hypothetical proteins SLIV_17115 and SLIV_17110 were tandemly encoded just upstream of the SLP structural proteins in the reverse direction (Fig. 1A; Supplemental Fig. 1). SLIV_17110 has a DUF4157 domain, which has been proposed to be the core domain of numerous CIS effector-related proteins (Fig. 1B) (20). Since interactions between a DUF4157 domain-containing protein and the CIS spike complex was suggested in a previous study on another *Streptomyces* CIS (21), we investigated the interaction between SLIV_17110 and predicted spike complex proteins SlpT2 (tube initiator; Afp5/Pvc5 homolog), Slp4 (spike hub; Afp7/Pvc7 homolog), and Slp5 (spike; Afp8/Pvc8 homolog) (Fig. 1A). His_6_-SLIV_17110 and FLAG-tagged spike complex protein were subjected to coelution assay using Ni^2+^-affinity chromatography, and potential interaction between SLIV_17110 and SlpT2 were detected (Fig. 1BC). The association between SlpT2 and SLP was confirmed by western blotting (Fig. 1D; Supplemental Fig. 2).

**Figure 1.**
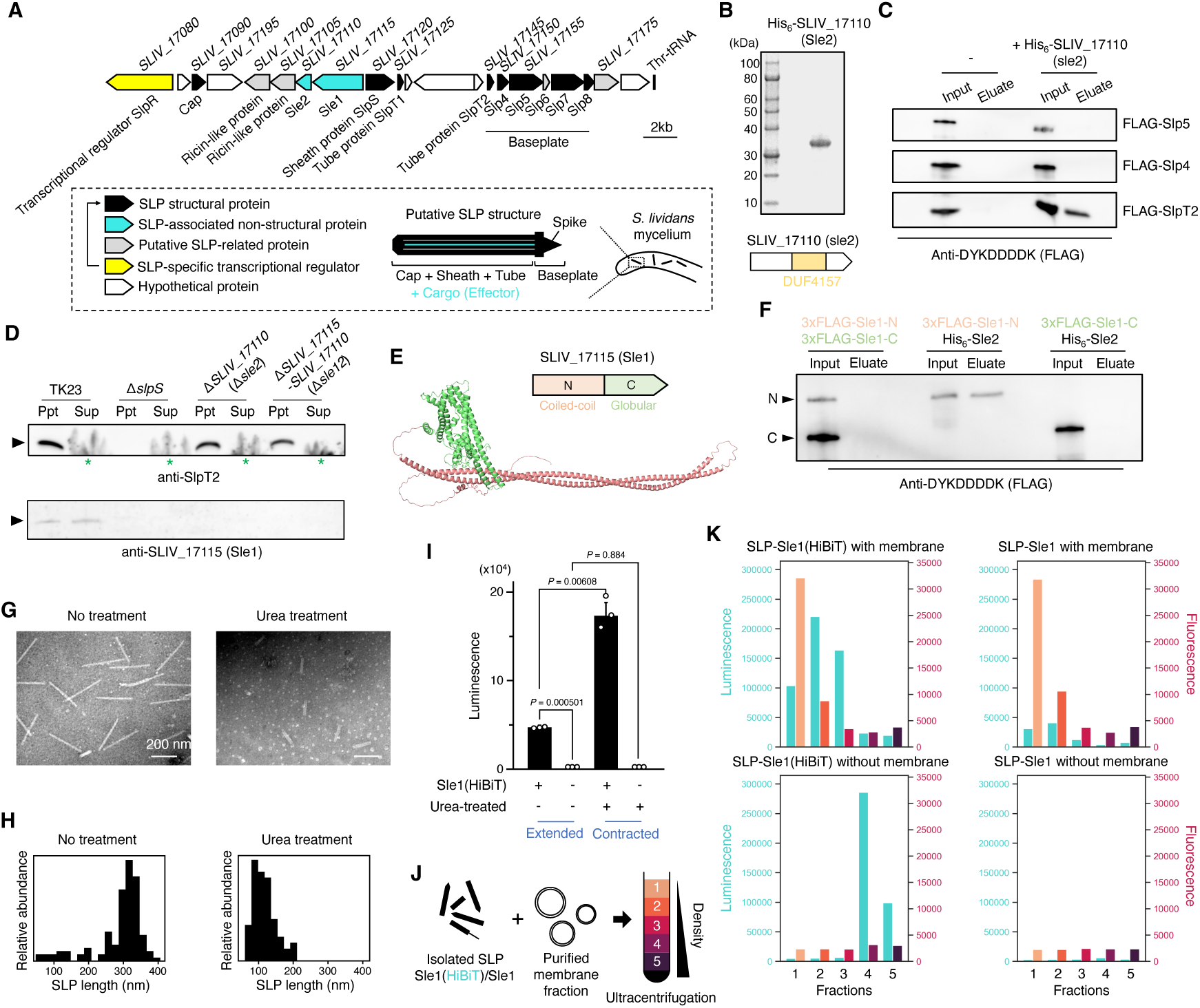
SLP loads a cargo protein that can be translocated to lipid membranes. An SLP cargo protein associated with a CIS core domain-containing protein was identified. (A) SLP gene cluster is shown. The SLP structural proteins were identified by proteome analysis and western blotting of the isolated SLPs. SlpR is a transcriptional regulator specific to the SLP gene cluster. A model of putative SLP structure is also shown inside a square with dashed lines. By similarity to CISs, a putative cargo protein (effector) is illustrated as being loaded inside the tube lumen of SLP. SLPs are localised in the cytoplasm of *S. lividans* mycelia. (B) SLIV_17110 containing the CIS core domain (DUF4157 domain) was cloned and purified as a hexahistidine (His_6_)-tagged protein. (C) DYKDDDDK (FLAG)-tagged SLP structural proteins SlpT2, Slp4, and Slp5 were individually mixed with purified His_6_-SLIV_17110 and then subjected to Ni^2+^-affinity chromatography. The inputs and eluates were analysed by western blotting using an anti-DYKDDDDK antibody. (D) Substrate mycelia of *S. lividans* grown on a solid medium were scraped off the plate and separated into SLP fractions (ppt; ultracentrifugation pellet) and detergent-soluble (sup; ultracentrifugation supernatant) fractions. These fractions were separated by SDS–PAGE and then subjected to western blot analysis. Green asterisks indicate that the SlpT2 signal was not clearly detected in the supernatant fractions because of the presence of a detergent, which would have interfered with the migration of SlpT2. (E and F) The structure of SLIV_17115 was predicted using AlphaFold3, and the predicted coiled-coil N-terminal and globular C-terminal regions were subjected to a co-elution assay with His_6_-SLIV_17110. DYKDHDGDYKDHDIDYKDDDDK (3×FLAG) was fused to the N-termini of the SLIV_17115 fragments. Tagged proteins were individually mixed with His_6_-SLIV_17110 and subjected to Ni^2+^-affinity chromatography. Western blotting was performed using the anti-DYKDDDDK antibody. (G) The isolated SLPs were treated with 3 M urea or buffer and observed by transmission electron microscopy after dilution to an appropriate concentration. (H) Length distributions of the images in *G* were analysed. Distributions of approximately 100 and 300 nm indicate the contracted and extended SLPs, respectively. (I) SLPs complemented with either SLIV_17115 (HiBiT) or native SLPs were isolated and treated with 3 M urea or buffer. After diluting to an appropriate concentration, the treated SLPs were subjected to the HiBiT-LgBiT split luciferase assay. Bars indicate mean ± S.D. for three independent assays. *P* values were calculated using *t*-test with Welch’s correction. (J) Experimental design of the interaction assay between SLP and lipid membranes. (K) Density-gradient ultracentrifugation samples were fractionated and subjected to the HiBiT-LgBiT split luciferase assay and FM1-43 staining. Luminescence and fluorescence indicate the presence of SLIV_17115 (Sle1) (HiBiT) and lipid membranes, respectively.

SLIV_17115, which is encoded just upstream of SLIV_17110, does not share close homology with any of the characterised proteins. This protein was predicted to consist of an N-terminal large coiled-coil segment and C-terminal alpha helices folded into a globular structure (Fig. 1E; Supplemental Fig. 3 and 4). Two of the C-terminal helices were predicted to form a transmembrane helical hairpin, showing remote homology with the membrane-anchor hairpin of the type III secretion system translocon component AopB (Supplemental Fig. 4) (22). The N-terminal region was predicted to have remote homology with the tapemeasure protein of the *Escherichia* phage P2 (Supplemental Fig. 5). Tapemeasure proteins are phage cargo proteins ejected from the tail tube lumen and have recently been proposed to be the evolutionary origin of some CIS effectors (21). Additionally, gene adjacency to DUF4157 domain-containing proteins has been recently proposed to be a characteristic feature of some CIS effectors lacking the DUF4157 domain (23). Since these observations imply the potential significance of SLIV_17115 as an effector associated with the CIS core domain-containing protein SLIV_17110, we performed an interaction assay for these proteins and found the interaction between SLIV_17110 and the N-terminal region of SLIV17115 (Fig. 1F). Furthermore, we identified 62 close SLIV_17115 homologs among 754 *Streptomyces* species with publicly available RefSeq genome sequences. Notably, all the SLIV_17115 homologs with the N-terminal extension were flanked by small DUF4157 domain-containing proteins and SlpS (sheath) homologs (Supplemental Fig. 6 and 7). Therefore, SLIV_17110 would be the conserved partner of SLIV_17115, and SLIV_17110 potentially mediates an indirect connection between SLIV_17115 and the baseplate core of SLP. Since the above results consistently suggested that these proteins are candidate SLP effectors, we named them SLP-associated effector 1 (Sle1) and Sle2, respectively.

We also performed a marker-less in-frame deletion of *sle1* and *sle2*. Although Δ*sle2* and Δ*sle1/2* mutants were obtained, our attempt to delete *sle1* alone failed. SLPs isolated from the Δ*Sle1/2* mutant did not show any apparent structural defects (Supplemental Fig. 8). However, the abundance of Sle1 was dramatically decreased by the deletion of *slpS*, which encode a sheath protein essential for SLP construction, and *sle2*, suggesting that mature SLP and Sle2 may enhance the biological stability of Sle1 (Fig. 1D; Supplemental Notes).

### An SLP effector SLIV_17115 (Sle1) is ejected upon SLP contraction and can be translocated to lipid membranes

To gain insights into the biological functions of the newly identified SLP-associated proteins, we first evaluated their effects on *E. coli* cells. We found that the expression of Sle1 markedly inhibited colony formation in *E. coli*, whereas Sle2 had no detectable effect, suggesting that Sle1 may be a cognate, biologically active effector of SLP (Supplemental Fig. 9). To investigate whether the contractile action of SLP externalises Sle1 (11,24), we explored the conditions that artificially triggered SLP contraction and found that exposure to urea efficiently induced SLP contraction, similar to the typical contractile phage tails (Fig. 1G and H) (25). We next introduced either Sle1-Sle2 or Sle1(HiBiT)-Sle2, with the latter construct including a 11-amino acids peptide tag (HiBiT tag) at the C-terminus of Sle1, into the Δ*sle1/2* mutant. The HiBiT tag activates a 19-kDa inactive luciferase fragment, LgBiT (26) (Supplemental Fig. 10). Upon the addition of LgBiT and a luminescent substrate (furimazine), significantly stronger luminescence was detected in the SLP fraction of the Sle1 (HiBiT)-Sle2-complemented strain (Fig. 1I). Notably, the luminescence of the SLP fraction of the Sle1(HiBiT)-Sle2-complemented strain markedly increased after exposure to urea, which induced SLP contraction (Fig. 1G-I). Thus, Sle1(HiBiT) would be loaded inside and ejected from the SLP during its construction being forced to be ejected by urea-induced sheath contraction. This allows more LgBiT, which is unlikely to permeate the extended SLP lumen, to access the activator HiBiT tag in Sle1(HiBiT).

Given the presence of the putative transmembrane region in Sle1 suggesting a potential association with the cellular membrane (Supplemental Fig. 4), we investigated the translocation of Sle1 to lipid membranes using the highly sensitive HiBiT-LgBiT system. Upon incubation of purified lipid membranes of the SLP-deficient (Δ*slpS*) mutant (16) and the isolated SLP containing Sle1(HiBiT), both the effector and SlpS migrated to the lipid membrane fractions (Fig. 1J and K; Supplemental Fig. 11). This result suggests that Sle1 can be translocated to lipid membranes along with SLPs.

### The C-terminal domain of Sle1 interacts with a ribosome-associated fraction and transiently delays the E. coli growth

To determine the Sle1 domain responsible for the inhibition of colony formation in *E. coli*, we assayed the truncated N- and C-terminal regions of Sle1. Only the C-terminal region (Sle1-C) exhibited an inhibitory effect on *E. coli* cells, indicating that this is a functional domain (Fig. 2A). Because some of effectors associated with phage tail-like nanomachines have been proposed to have bactericidal activities (20), we investigated whether Sle1-C kills *E. coli* by measuring growth of the *E. coli* strains in liquid medium with or without the induction of Sle1-C. Notably, Sle1-C expression resulted in a significant delay in growth, and Sle1-C-expressing *E. coli* cells in the stationary phase were resuscitated immediately after inoculating into fresh medium without the inducer (Fig. 2B and C). Therefore, Sle1-C is likely to cause a transient delay in *E. coli* growth rather than killing the bacterium through bactericidal activity.

**Figure 2.**
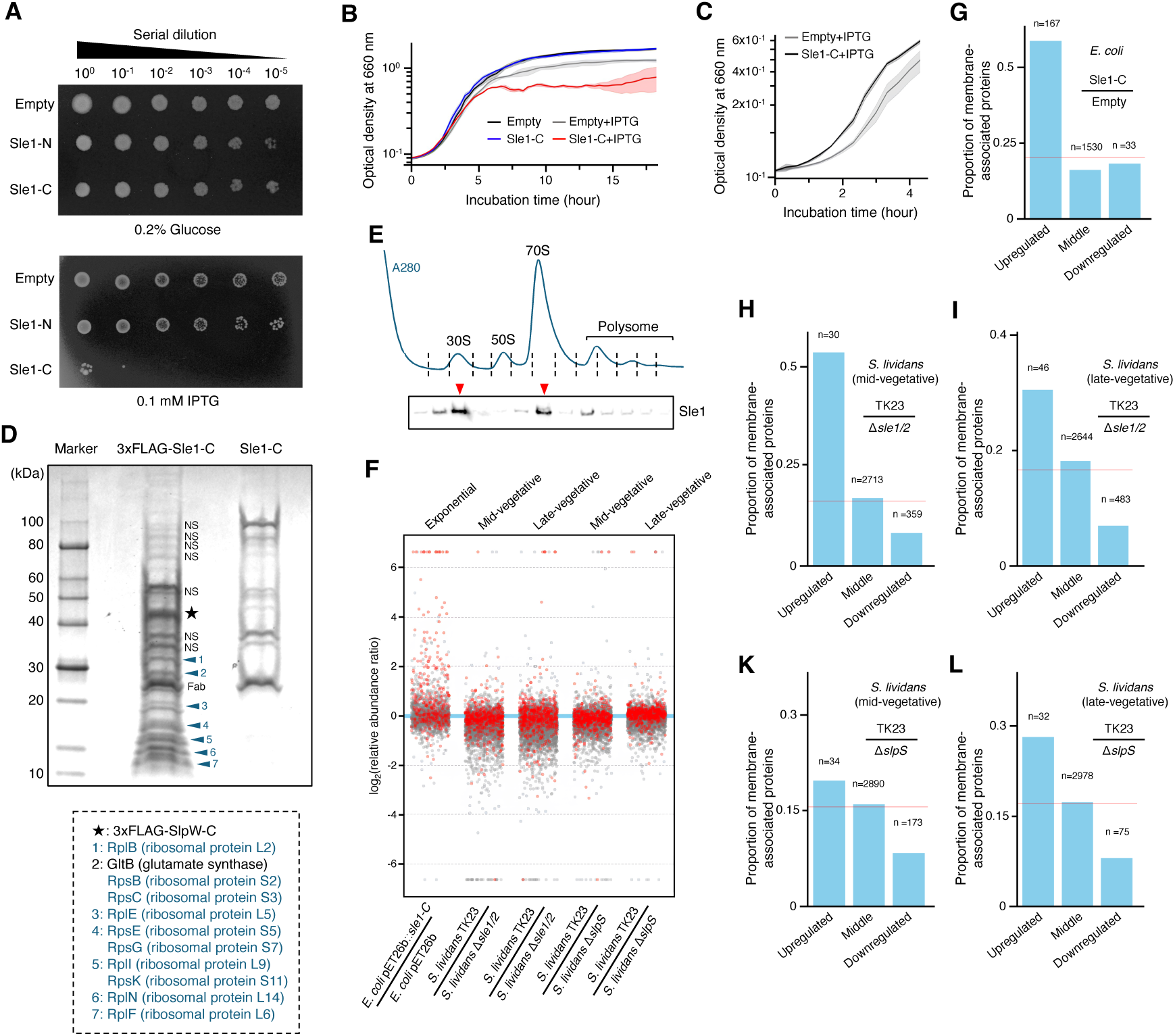
Characterisation of Sle1 as an SLP effector influencing cellular membrane-associated bacterial proteome. SLIV_17115 (Sle1) is characterised as an SLP effector that affects the protein profiles in both *E. coli* and *S. lividans*. (A) Cultures of *E. coli* NiCo21(DE3) harbouring pET26b, pET26b::*sle1-N*, or pET26b::*sle1-C* were serially diluted and spotted onto solid LB medium containing a suppressor (0.2% [w/v] glucose) or inducer (IPTG, 0.1 mM). (B) Growth curves of *E. coli* NiCo21(DE3) cells harbouring pET26b or pET26b::*sle1-C* are shown. IPTG (0.025 mM) was added as the inducer. Lines and coloured areas indicate the mean values ± S.D., respectively, for three independent cultures. (C) The *E. coli* strains used in panel *B* were grown to the stationary phase in the presence of IPTG and these cells were then regrown after diluting to the same optical density at 660 nm in fresh liquid LB medium. Lines and coloured areas indicate the mean values and S.D., respectively, for three independent cultures. (D) *E. coli* NiCo21(DE3) cells harbouring pET26b::*sle1-C* or pET26b:: *3×FLAG -sle1-C* were grown to the mid-exponential phase in the presence of the inducer and then 3×FLAG-Sle1-C was pulled down using anti-DYKDDDDK antibody fragment (Fab)-conjugated agarose beads. Potential interacting proteins were identified using MALDI-TOF-MS and MASCOT search. NS, non-specific band. (E) Ribosomes were extracted from the substrate mycelia of *S. lividans* and purified by sucrose density-gradient ultracentrifugation. Each fraction was subjected to western blot analysis using anti-Sle1 serum. (F) Total proteins were extracted from the indicated bacterial strains and subjected to quantitative proteome analysis. The abundance ratio refers to the abundance of each detected protein (scatter plot) in the reference strain (numerator) relative to that in the control strain (denominator). Proteins with inferred membrane localisation are shown in red. The upper and lower limits for calculating of the abundance ratio were 100- and –100-fold, respectively. (G-L) Proportions of the predicted membrane proteins detected by the proteome analysis in panel *F* were calculated for the following three categories: upregulated, abundance ratio ≥ 2; middle, –2 < abundance ratio < 2; downregulated, abundance ratio ≤ –2. Red lines indicate the mean proportions of the predicted membrane proteins among the total proteins commonly detected in the two compared strains.

We could not find any significant structural motifs with known enzymatic functions in Sle1-C, leading to the hypothesis that Sle1-C delays *E. coli* growth by a mechanism other than enzymatic disruption of biomolecules. Based on this hypothesis, we performed a co-immunoprecipitation assay to identify Sle1-C targets in *E. coli*. 3×FLAG-tagged Sle1-C or the non-tagged version was heterologously expressed in *E. coli* and the cell lysates were mixed individually with agarose beads conjugated with anti-DYKDDDDK (FLAG tag) antibody fragments. Subsequent mass spectrometry analysis identified most of the proteins associated with the immunoprecipitated Sle1-C as ribosomal proteins constituting the 30S and 50S subunits of the *E. coli* ribosome (Fig. 2D). Given the potential association of heterologously expressed Sle1-C with ribosomal proteins, we also analysed the ribosome fractions isolated from *S. lividans* mycelia and found that Sle1 was mainly detected in the 30S and 70S ribosome fractions, indicating an association between Sle1 and the subcellular fractions containing the small ribosomal subunit in *S. lividans* (Fig. 2E).

### Sle1 upregulates membrane protein-rich sub-proteome in both E. coli and S. lividans

Since Sle1-C did not inhibit the activity of the protein synthesis system in *E. coli* (Supplemental Fig 12; Supplemental Notes), we assumed that Sle1-C might indirectly alter the protein expression profiles of *E. coli* cells by interacting with the ribosome-associated fraction rather than inhibiting ribosomal activity. To test this hypothesis, we performed quantitative proteome analysis of the *E. coli* strains harbouring either the Sle1-C expressing plasmid or the empty plasmid. Overall, at the mid-late stage of the exponential growth phase, Sle1-C expressing cells were more abundant in experimentally or bioinformatically identified membrane proteins than in control cells (Fig. 2F and G; Supplemental Data). Statistically significant correlations between the abundance ratios relative to the control sample and the annotations as membrane proteins were only detected in the two proteome subsets with abundance ratios of ≥2.0 (167 proteins; point-biserial correlation coefficient, 0.243; *P* value, 0.00158) or ≥1.0 and <2.0 (698 proteins; point-biserial correlation coefficient, 0.0865; *P* value, 0.0222). The proteome subset with an abundance ratio of ≥2.0 included membrane proteins that play roles in respiration (NADH-ubiquinone oxidoreductase subunits A, N, H, and L), saccharide uptake (PTS system EII components), peptidoglycan synthesis (MurJ), and protein export (SecD/F/Y, tatE). Thus, it is possible that over-accumulation of these membrane proteins may cause imbalance of metabolic fluxes, limiting the ability of the cells to adapt to varying nutritional conditions and consequently leading to slower growth. Furthermore, western blot analysis using purified lipid membranes from *E. coli* indicated that Sle1-C was partially localised to the membranes, consistent with the functional relevance of Sle1-C to membrane proteins (Supplemental Fig. 13).

Given the significant changes in the *E. coli* proteome upon Sle1-C expression, we further performed proteome analysis of the *S. lividans* strains TK23, Δ*slpS*, and Δ*sle1/2*. The results revealed an overall downshift in the relative protein abundance ratios in the parental TK23 strain compared to the deletion mutants during the mid-late vegetative growth phase (Fig. 2F). No consistent upregulation of stress response-related proteins, such as major molecular chaperones and heat shock proteins, was observed (Supplemental Fig. 14; Supplemental Data). In contrast, we also found that experimentally or bioinformatically identified membrane proteins were consistently abundant in the proteome subsets where the relative abundance ratios of proteins were significantly higher (> 2-fold) in the parental TK23 strain than in either of the deletion mutants. The opposite trend was observed in the subsets where the abundance ratios were lower in the TK23 strain (Fig. 2F and H-L; Supplemental Data). Although we did not observe molecular species-specific changes in protein abundance in the above proteome subsets across strains and time points, the NADH-ubiquinone oxidoreductase subunit E was significantly less abundant in both the Δ*slpS* (0.461 relative to TK23) and Δ*sle1/2* (0.373 relative to TK23) at the mid-vegetative growth stage, implying a potential effect on energy metabolism (Supplemental Data).

### The Sle1/2 pair tunes the metabolic activity of S. lividans to maintain reproductive activity under competition

Given the broad impact of Sle1 on the *S. lividans* proteome, we were interested in how this was reflected in the phenotypes of this bacterium. We measured the vegetative growth of the parental TK23 strain and the Δ*sle1/2* mutant and obtained similar growth curves (Fig. 3A). Spore formation following the vegetative growth of substrate mycelia in the *Streptomyces* life cycle (27) was also comparable between these strains (Fig. 3B). We next investigated energy metabolism of the *S. lividans* strains, since Sle1-associated enrichment of energy metabolism-related membrane proteins were consistently observed (Fig. 2F-L; Supplemental Data). We found that substrate mycelia of the Δ*sle1/2* mutant showed lower reducing activity than the parental TK23 strain and the *sle1/2*-complemented strain (Δ*sle1/2 attC*::*sle1/2*) at the mid-late stage of vegetative growth, indicating that loss of the Sle1/2 pair would lead to lower energy metabolism at this stage (Fig. 3C). We also examined the capacity of the substrate mycelia for energy-consuming protein synthesis by monitoring inducible msfGFP expression in *S. lividans* strains. Notably, the linear regression slope of msfGFP fluorescence of the Δ*sle1/2* mutant was lower than that of the TK23 strain at the mid-late stage of vegetative growth, which is consistent with the lower energy metabolism of the mutant (28) (Fig. 3D). In addition, the involvement of SLPs in the above differences was also suggested since the Δ*slpS* mutant also showed lower cellular reducing activity and msfGFP induction than the parental TK23 strain at the same growth stage (Fig. 3C and D). In contrast, at the early stage of vegetative growth, the Δ*sle1/2* mutant showed higher cellular reducing activity and msfGFP induction (Supplemental Fig. 15). These results indicate that the Sle1/2 pair may repress the consumption of resources independently of SLPs at the early stage of vegetative growth, whereas Sle1/2 and SLP may co-act to maintain the metabolic capacity at the later stages.

**Figure 3.**
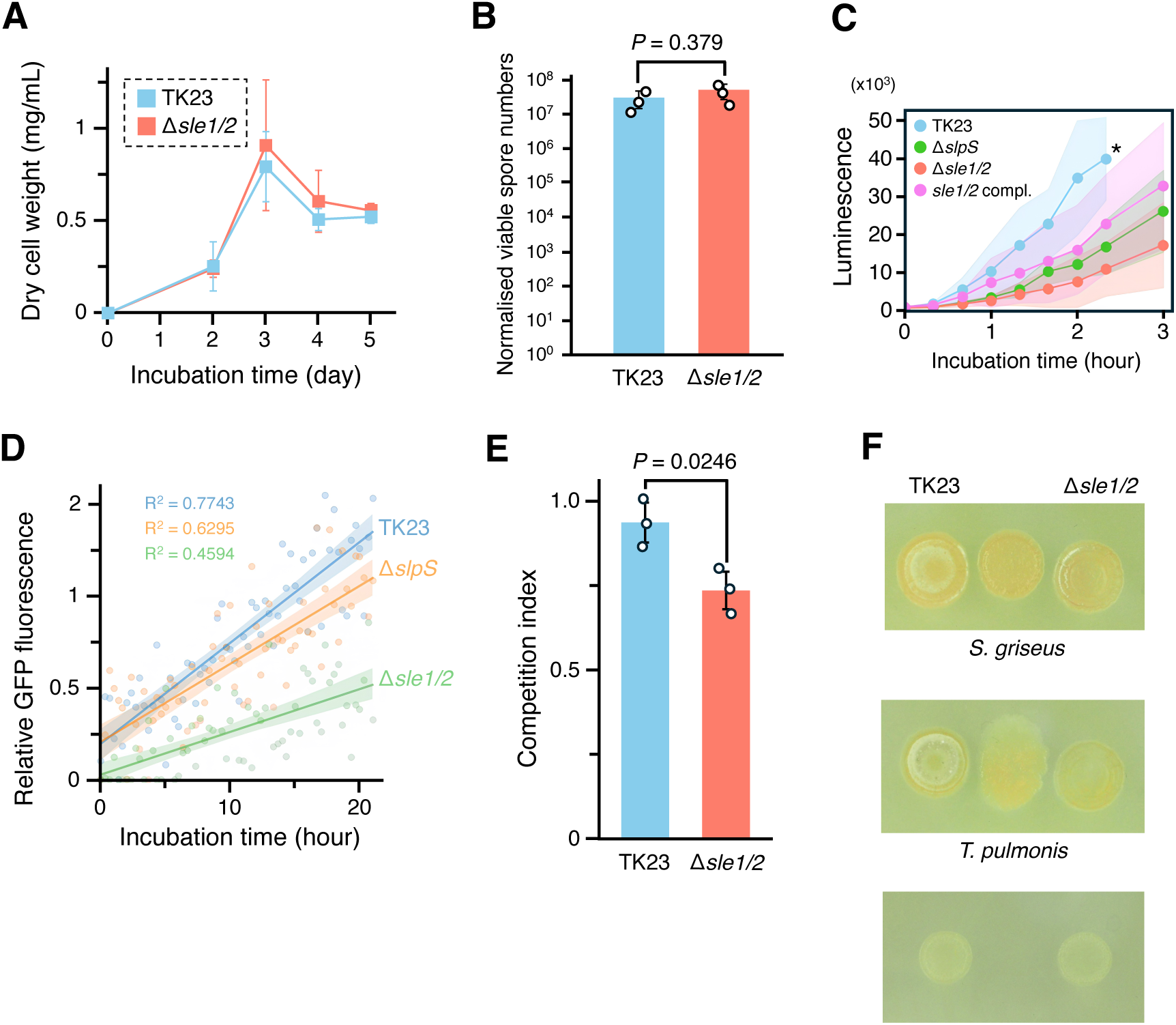
The Sle12 pair contributes to the maintenance of reproductive activity of *S. lividans* under competition. The Sle12 pair potentially confers an ecological benefit to *S. lividans* by enhancing its adaptability under competition for nutrients. (A and B) Vegetative growth and spore formation were compared between the parental TK23 strain and the Δ*sle12* mutant by measuring dry cell weight in liquid media and the number of viable spores formed on solid media, respectively. Values and error bars indicate means ± S.D. for (A) five or (B) three independent cultures. (C) The cellular reducing activity of each *S. lividans* strain was measured using a cell-permeable luminescent substrate and luciferase as described in the Methods section. Values and coloured areas indicate means ± S.D. for three independent cultures. The asterisk indicates the time point after which the luminescence of ‘TK23’ exceeded the detection threshold. (D) Protein synthesis capacity of *S. lividans* strains expressing msfGFP was measured. In these constructs, msfGFP expression was regulated by an engineered *tipA* promoter (P*_tipA_*RS), in which the transcription and translation of the downstream open reading frames were dependent on exogenously added thiostrepton and theophylline, respectively. Relative GFP fluorescence was calculated by subtracting the fluorescence of mycelia under non-inducing conditions from that of mycelia under inducing conditions. The *S. lividans* strains in this panel represent the genetic background of the msfGFP-expressing strains. Scatter plots, lines, and coloured areas indicate the mean values of three independent samples, regression lines, and 95% confidence intervals, respectively. (C) *S. lividans* strains harbouring a thiostrepton resistance gene and *Streptomyces griseus* were inoculated on a solid sporulating medium and spores were collected from the co-culture. The competition index was calculated as the viable spore number of *S. lividans* relative to that of *S. griseus*. The *S. lividans* strain in this panel represents the genetic background of the thiostrepton-resistant strain used for the competition assay. (F) *S. lividans* strains were co-cultured with *S. griseus* or *Tsukamurella pulmonis* on solid media without direct contact for three days. The white regions of the colonies indicate aerial mycelia.

Since the decrease in energy metabolism and protein synthesis in the Δ*sle1/2* mutant at the later growth stage suggests the vulnerability of this mutant to a nutrient-limited condition, we tested various growth conditions under which *S. lividans* potentially experiences low-nutrient stress. We found that the spore formation of this strain was relatively lower than that of the parental strain in co-culture with *Streptomyces griseus*, which competes with *S. lividans* for nutrients (Fig. 3E; Supplemental Fig. 16). In addition, we observed an apparent delay in aerial mycelia erection in the Δ*sle1/2* mutant colony co-cultured with the competitor without direct contact (Fig. 3F). A delay in mycelial erection was also observed in the mutant colony in a co-culture with *Tsukamurella pulmonis*, which has been characterised as intimately interacting with *Streptomyces* species in nature and not producing antibiotics significantly effective against *S. lividans* (29–31) (Fig. 3F).

### A wide distribution of Sle1/2-like proteins in the major actinobacterial class and their functional diversity

Given the unique function of Sle1, we were interested in whether Sle1 and its partner Sle2 are conserved among bacteria. We first searched for Sle2 homologs using the eCIStem database (20) and found that they were located within various CIS-related gene clusters conserved in the phylum Actinobacteria (Fig. 4). Notably, they were frequently encoded adjacent to alpha helix-rich proteins and just upstream of CIS sheath protein homologs. The alpha helix-rich proteins include large coiled-coil segments at the N-termini, and share remote homology with phage tapemeasure proteins (Supplemental Fig. 17). All these structural features and genomic contexts are consistent with those of Sle1, suggesting that Sle1/2-like pairs are widely shared across actinobacteria, probably serving as cognate effectors for CIS-related nanomachines beyond *Streptomyces*.

**Figure 4.**
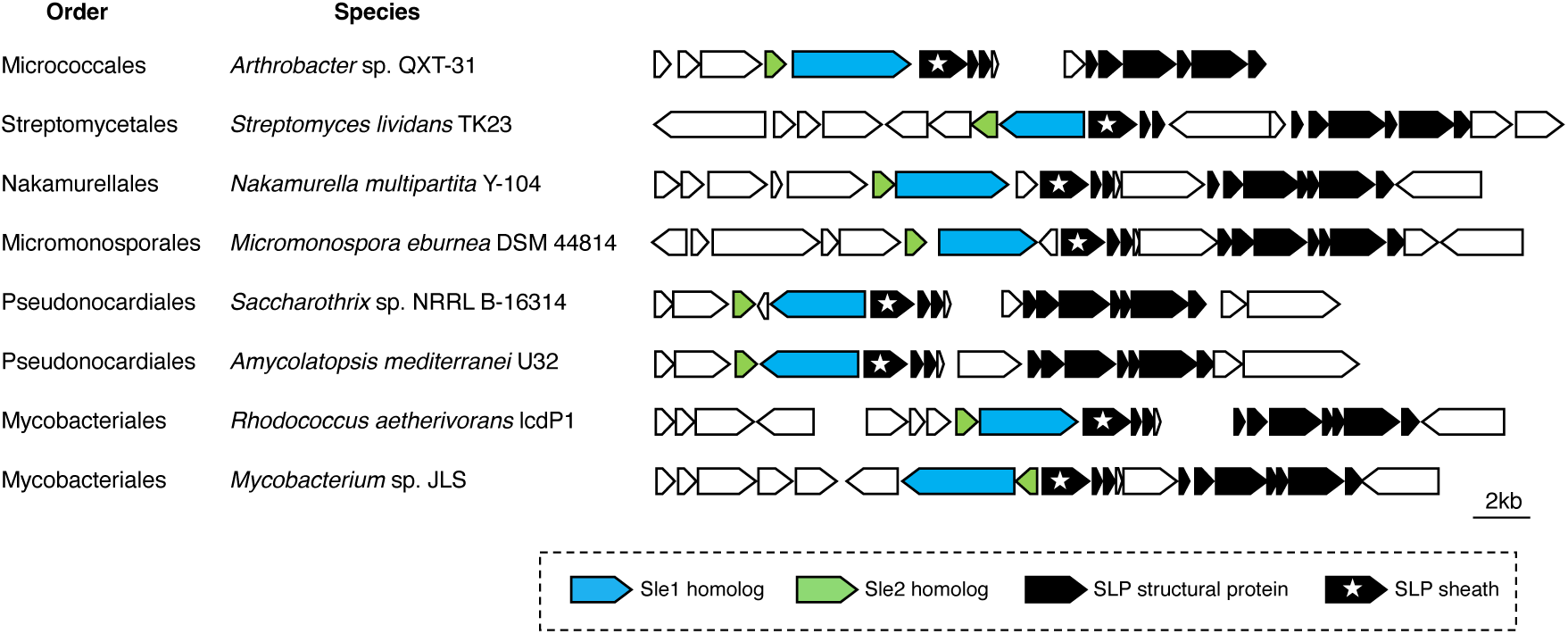
Distribution of Sle12-like pairs in the major actinobacterial class. Homologs of Sle12 are widely conserved in the class Actinobacteria with consistent gene synteny.

Importantly, the C-terminal regions of the identified Sle1-type proteins showed little similarity to each other, implying the functional diversification of the putative Sle1-type effectors. To examine this possibility, we cloned the C-termini of Sle1-type proteins RS14790 from *Micromonospora eburnea* DSM 44814 and RS24535 from *Amycolatopsis mediterranei* U32. Whereas RS24535 showed no detectable effect, RS14790 significantly inhibited *E. coli* growth presumably by its enzymatic activity on nucleic acids (Fig. 5A; Supplemental Fig. 18 and 19). Furthermore, we also found that a hypothetical protein RS14785 encoded just downstream of RS14790 significantly alleviated the growth inhibition caused by RS14790 upon co-expression (Fig. 5A-C). Thus, these proteins would constitute an effector-immunity system associated with an SLP-like nanomachine. Notably, an Sle1-associated immunity protein was not found in the SLP gene cluster, suggesting that the RS14790/RS14785 system is functionally distinct from Sle1, although it retains the key features of Sle1-type effectors.

**Figure 5.**
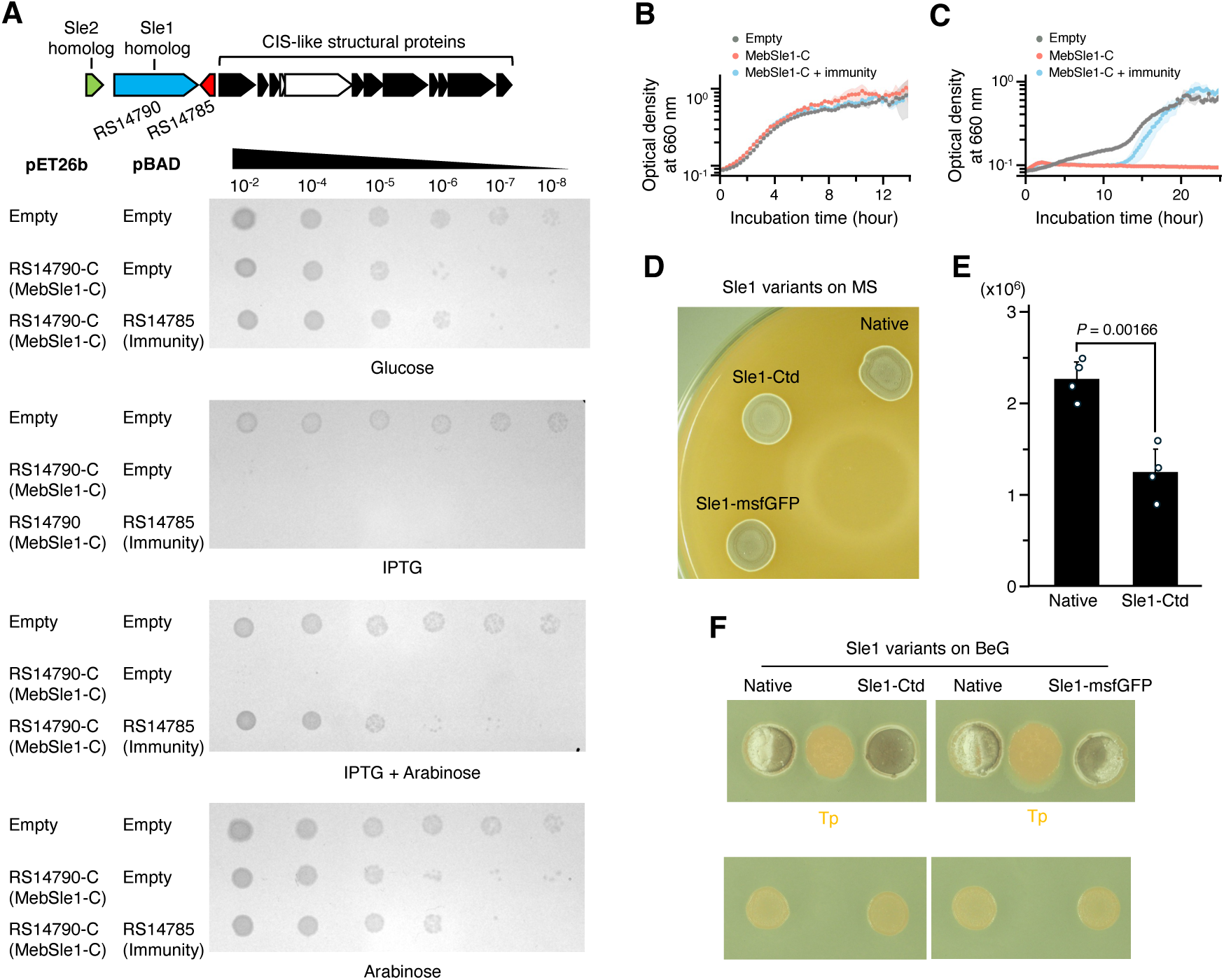
Functional diversification and reconfigurability of Sle1-type effectors. The functional domain of the Sle1-like protein associated with the CIS-related gene cluster of *M. eburnea* was identified, and the effect of appending this domain to Sle1 was investigated. (A) *E. coli* strains harbouring either of the pET26b/pBAD plasmid sets were cultured, serially diluted, and spotted onto solid LB medium containing 50 μg/mL antibiotics and additives (0.2% [w/v] glucose as a suppressor; 0.15 mM IPTG as an inducer for pET26b; 0.2%[w/v] arabinose as an inducer for pBAD. (B) The *E. coli* strains used in panel *A* were cultivated in liquid LB-Lennox media containing kanamycin and ampicillin. 0.5% (w/v) Glucose and arabinose were added to the media as the suppressor and inducer for pET26b and pBAD, respectively. (C) Glucose in the cultures in panel *B* was replaced with 0.25 mM IPTG to induce the protein expression of pET26b. (D) 10^4^ viable spores of each strain were spotted onto solid MS medium and incubated for 4 days. Note that the spore maturation of *S. lividans* colonies involves the colony colour turning from white into grey. (E) Spores were collected from colonies grown on solid MS medium and then the number of viable spores was measured. Values and error bars indicate means ± S.D. for four independent cultures. *P* values were calculated using *t*-test with Welch’s correction. (F) *S. lividans* strains were co-cultured with *T. pulmonis* on a solid medium without direct contact for 5 days. The white and grey regions of the colonies indicate aerial mycelia and subsequent spore maturation, respectively.

### Sle1 can alter the phenotypic response of S. lividans to neighbouring bacteria by acquiring an exogenous functional module

To test whether functional diversification of Sle1-type effectors influences phenotypes of a producer bacterium, we fused the cytidine deaminase-like domain (Ctd) of RS14790 from *M. eburnea* with the C-terminus of Sle1 and introduced the resultant artificial Sle1 variant, Sle1-Ctd, into *S. lividans*. The strain possessing Sle1-Ctd formed a slightly whitish colonies on the spore formation medium (MS medium) compared to the control strains possessing either the native Sle1 or the msfGFP-fused Sle1, suggesting limited spore maturation (Fig. 5D) (32,33). Consistent with this observation, formation of viable spores was significantly lower in the strain posessing Sle1-Ctd than in the control strain (Fig. 5E). Since these strains showed no other apparent phenotypic differences in monoculture, we tested co-culture with *T. pulmonis*, which promoted visible phenotypes associated with Sle1 (Fig. 3). In contrast to the phenotypes observed in the monoculture, the strain expressing Sle1-Ctd formed grey regions on the colony more rapidly upon approaching than the control strains, indicating faster formation of mature spores (Fig. 5F; Supplemental Fig. 20). These results suggest that functional modification of Sle1 can alter the phenotypic response of *S. lividans* to neighbouring bacteria.

## Discussion

A series of data suggests that Sle1, encoded just upstream of the sheath protein SlpS, is a cargo loaded into SLP. Sle1 contains the coiled-coil region spanning 440 amino acids at the N-terminus, which is physically and genetically associated with the DUF4157 domain-containing proteins and shows remote homology with the coiled-coil regions of phage tapemeasure proteins. Although the detailed loading mechanism of tapemeasure proteins is not fully understood, several cryo-EM studies have shown that the coiled-coil segments of tapemeasure proteins interacting with the spike-baseplate complex extend along the lumen and are surrounded by organised tube protein complexes (34–37). Tapemeasure proteins are ejected upon sheath contraction of tailed phages and are then inserted into the cellular membrane, thereby playing a role in the transfer of viral genetic material into target cells (38–41). It is noteworthy that the tapemeasure protein-related effector (Tme) in *Streptomyces davawensis* shares the basic action mechanism with phage tapemeasure proteins, suggesting an evolutionary relationship between certain CIS effectors and the phage cargo (21). The findings of our study would expand the generality of this concept to a broader range of CIS effectors, including Sle1. However, the lack of the CIS core domain DUF4157 was a significant difference distinguishing Sle1 from Tme-like effectors, implying distinct molecular mechanisms underlying their associations with SLP/CIS. Thus, Sle1 would represent a novel class of tapemeasure protein-like CIS effectors, providing the new definition of CIS effectors (13,14,20,23).

As recently demonstrated in a tapemeasure protein of another phage tail-like nanomachine (37), the predicted globular C-terminal domain of Sle1, which consists of alpha-helices connected by turns/loops, may adapt its conformation to the tube lumen before ejection (Fig. 1E). Upon sheath contraction, Sle1 is ejected from the particle, and then the C-terminal domain would be relieved from the spatial constraints of the tube lumen and fold into a globular structure. Given the presence of the predicted transmembrane helices in the C-terminal domain of Sle1 and the proposed membrane insertion mechanism of its remote homolog AopB (Supplemental Fig. 4) (22), Sle1 would be translocated to the cellular membrane, and SLP may facilitate this process (Fig. 1J and K). Regarding this possibility, it should be noted that the closely related CIS*^Sc^* might transiently interact with a putative membrane-associated protein which possibly induces sheath contraction and membrane insertion of an effector(s) (42). Because the putative membrane-associated protein (SLIV_17175, Fig. 1A) is also conserved in the SLP gene cluster, Sle1 may be translocated to the cellular membrane through a similar mechanism (Fig. 1A). Furthermore, the potential interactions with ribosomal proteins would suggest that Sle1 might act as a molecular hub linking the protein synthesis machinery and cellular membrane, thereby facilitating the expression of membrane proteins transcribed in the proximity of Sle1, although this functional model would need further verification (Fig. 2F-L). Tapemeasure proteins of some actinobacterial phages have long been known to contain C-terminal motifs that potentially modulate host physiology for phage reproduction, implying their alternative roles as viral signaling molecules (43–45). Thus, it is possible that the remarkable tolerance of phage tapemasure proteins to structural variations might have allowed for the emergence of Sle1-type effectors with the unique biological functions.

The *Streptomyces* life cycle encompasses spore germination and substrate mycelial extension along and beneath the surface of the medium, eventually turning into aerial mycelia erecting into the air. The transition from vegetative growth of substrate mycelia to aerial mycelial erection often occurs in response to external nutritional deprivation (27). Therefore, this unique process has to largely rely on the reuse of cellular building blocks and the consumption of energy sources stored within the biomass of substrate mycelia at the late stage of vegetative growth (46). Aerial mycelia finally form unicellular spores that are more persistent under nutrient limitations and are crucial for increasing the overall fitness of *Streptomyces* colonies (47). Therefore, active substrate mycelial biomass would be one of the factors determining colony fitness, and its importance in *Streptomyces* ecology could be more significant in natural settings, where microorganisms compete spatially for limited nutrition. From this perspective, our findings on Sle1 may suggest its possible role in maintaining active biomass containing building blocks and energy sources that can be readily utilised for spore formation under competitive conditions. Sle1 would direct protein synthesis to the relatively small membrane-associated proteome subset while decreasing the larger subset rich in cytoplasmic proteins, possibly avoiding excessive resource consumption (Fig. 2 H-L). Among the membrane proteins upregulated in the parental strain, those crucial for aerobic respiration would be notable since their reduction in the relative abundance may explain the decreased metabolic activity of the Δ*slpS* and Δ*sle1/2* mutants (Fig. 3; Supplemental Data). Although functions of the upregulated hypothetical membrane proteins remain unclear, we speculate that they might contribute to the proper localisation and/or stabilisation of other membrane proteins, including those essential for energy metabolism and the maintenance of cell envelope integrity. Additionally, the robust regulatory linkages between SLP production and nutritional and cell envelope stress (16,17,19,48) support the idea that the significance of SLP and its effector Sle1 is relevant to the fitness of *S. lividans* under ecological stress, which could compensate for the high metabolic cost of producing this nanomachine (49).

Our results also have ecological implications of the diversification of Sle1-type effectors among actinobacterial species. We show that the fusion of Sle1 with the deaminase-like domain led to faster spore maturation in response to neighbouring bacteria, indicating that Sle1-type effectors can affect producer phenotypes under competition by acquiring a new functional module at the C-terminus. Although it remains unclear how the deaminase-like domain alters the pattern of spore formation, we assume that moderate genotoxic stress caused by this domain may activate stress response pathways and render *S. lividans* more prepared to proceed with sporulation in response to microbial competition. Such mutation in Sle1-type effectors could be beneficial under certain ecological circumstances, eventually leading to the conservation of functional motifs/domains in these effectors and ultimately their diversification. The evolutionary selection of CIS effectors may also explain the variations in CIS-associated phenotypes among *Streptomyces* species (Supplemental Notes).

Taken together, the present study identified the novel group of effectors associated with phage tail-like nanomachines that are highly conserved in actinobacterial species. The inferred evolutionary background of these effectors suggests that, in the end of ancient host-virus arms race, the potential of the phage infection machinery have been unleashed and harnessed as a reconfigurable tuner for cellular functions. Our findings shed light on the previously untapped resource of CIS effectors and connect them to ecological trait modulation, expanding the applicability of the CIS repertoire as versatile bio-tools.

## Methods

### Culture conditions

Strains used in this study are listed in Supplemental Table 1. *M. eburnea* NBRC 101912 and *A. mediterranei* NBRC 13415 were obtained from NBRC (Chiba, Japan). *Streptomyces lividans* TK23 and *S. griseus* IFO13350 were routinely grown in Bennett’s-glucose medium comprising 0.1 g yeast extract, 0.1 g meat extract, 0.2 g N-Z amine, 1 g glucose per 100 mL (pH7.2). For solid media, 2% (w/v) agar was added. For spore formation, *Streptomyces* species were grown on mannitol-soya flour (MS) medium comprising 2 g mannitol, 2 g soya flour, and 2 g agar per 1 L. For extraction of SLP and lipid membranes, *S. lividans* spores (approximately 10^4^ viable spores) were inoculated on a sterilised cellophane membrane placed onto Bennett’s-glucose medium and then the colonies were scraped off the membrane for further treatments. For growth measurements, *S. lividans* was precultured in 4 mL Bennett’s-glucose medium and then the preculture was inoculated into 100 mL YPD medium comprising 1 g yeast extract, 2 g peptone, 2 g glucose per 100 mL. Note that YPD medium was used to obtain reproducible growth curves of this bacterium. *T. pulmonis* was grown in liquid tryptic soy broth medium for preculturing. *E. coli* strains were grown in LB-Lennox medium comprising 5 g yeast extract, 10 g tryptone, and 5 g NaCl per 1 L.

### Genetic manipulations

Primers and plasmids used in this study are listed in Supplemental Table 2. NEBuilder (New England Biolabs, MA, USA) and In-Fusion (Takara Bio Inc., Shiga, Japan) were used for sequence-independent DNA cloning. KOD One (Takara Bio Inc.), Prime STAR Max (Takara Bio Inc.), and Ex Premier (Takara Bio Inc.) were used for PCR.

For marker-less in-frame gene deletion, flaking regions of *SLIV_17110 or SLIV_17110-SLIV_17115* were amplified by PCR using primer sets (Sle2-FR1_Fw/Rv; Sle2-FR2_Fw/Rv; Sle1/2-FR2_Fw_Rv) and fused with pK18mob plasmid digested with EcoRI and HindIII. The resultant plasmids (pk18mob::*sle2*-FR12 and pk18mob::*sle1/2*-FR12) were amplified in *E. coli* DH5α and then each of the plasmids was introduced into *E. coli* S17-1. The transformed *E. coli* was grown in liquid LB-Lennox medium containing 50 μg/mL kanamycin and 50 μg/mL streptomycin, and the cells at the early exponential growth phase were collected from 1 mL of the culture. The collected *E. coli* cells were resuspended with *S. lividans* spore solution (approximately 10^7^ viable spores) and the mixture was spread onto a MS medium. After incubation at 30°C for 16-20 h, kanamycin and aztreonam were added to the culture to the final concentrations of 20 μg/mL. The transconjugants appeared after additional incubation for 2-3 days were isolated and streaked onto a fresh Bennet’s-glucose medium containing kanamycin and aztreonam. The sub-cultured colonies were grown on a solid MS medium for spore formation and the spores were spread onto solid Bennett’s-glucose media with dilution to approximately 100 spores per plate. The deletion mutants were selected from the resultant colonies by a kanamycin sensitivity assay and colony PCR.

For integration of gene cassettes into the *S. lividans* chromosome, integration plasmids were constructed as follows. For the construction of the *Sle1/2* complemented strains, *Sle1-Sle2* and *Sle1-hibit* sequences including the native promoter were amplified using primer sets (Sle1/2_pTYM19t_Fw, Sle1/2_pTYM19t_Rv, Sle1(HiBiT)_Rv). For the construction of the HiBiT tag-fused Sle1, *Sle2* sequence was separately amplified using primer set (HiBiT-Sle2_Fw, Sle1/2_pTYM19t_Rv). Each of *Sle1-Sle2* and *Sle1-hibit*/*Sle2* fragments was fused with pTYM19t (50) plasmid digested with KpnI and HindIII. For *Sle1-hibit/Sle2*, a hybridised linker sequence (GGGGSx2-HiBiT_Fw/Rv) was also added to the reaction. For the construction of the msfGFP-expressing strains, P*_tipA_*-RS, in which the ribosome binding site of the *tipA* promoter is replaced with a theophylline-inducible riboswitch (21), and codon-optimised *msfgfp* were fused with pTYM19t as described above. Each pTYM19t-derived plasmid was amplified in *E. coli* DH5α and introduced into *E. coli* S17-1 for the conjugation. Transconjugants were selected using thiostrepton. For the structural modification of Sle1, *msfgfp*, *RS14790-C*, *sle1*, and *sle2* sequences were amplified using primer sets (msfGFP_Sle1_Fw/Rv; RS14790-C_Sle1_Fw/Rv; Sle1/2_pTYM19t_Fw, Sle1(HiBiT)_Rv; Sle2_Fw, Sle1/2_pTYM19t_Rv) and fused with the linker sequence and the digested pTYM19t plasmid as described above.

For heterologous protein expression in *E. coli*, plasmids were constructed as follows. For the expression using pET26b plasmid, insert sequences were amplified by PCR using primer sets (SlpT2_Fw/Rv; Slp4_Fw/Rv; Slp5_Fw/Rv; Sle1-N_Fw/Rv; Sle1-C_Fw/Rv; RS24535-C_Fw/Rv; RS14790-C_Fw/Rv) and then fused with the plasmid digested with NdeI and HindIII. If necessary, nucleic acid sequences encoding DYKDDDDK (FLAG) or DYKDHDGDYKDHDIDYKDDDDK (3×FLAG) were fused with the insert sequences and the digested plasmid. For the expression using pET15b, insert sequences were amplified by PCR using primer sets (Sle1_pET15b_Fw/Rv; Sle2_pET15b_Fw/Rv) and then fused with the plasmid digested with NcoI and BamHI. For the expression using pColdII, insert sequences were amplified by PCR using primer set (Sle2_pCold_Fw/Rv) and then fused with the plasmid digested with BamHI and HindIII. For the expression using pMAL-c6T, the codon-optimised *sle1-C* sequence was amplified by PCR using a primer set (Sle1-C_pMAL_Fw/Rv) and then fused with the plasmid digested with NotI and HindIII. For the expression using pBAD/His A, *RS14785* sequence were amplified by PCR using primer set (RS14785_Fw/Rv) and then fused with pBAD/His A digested with NcoI and HindIII.

### Extraction of SLPs and ejection assay

SLPs were extracted from *S. lividans* mycelia as follows. Spore solutions (approximately 10^4^ viable spores) were spread onto solid Bennett’s-glucose media. After incubation at 30°C for 2 days, the colonies were scraped off the plate and then lysed in the solution comprising 20 mM HEPES-NaOH (pH7.5), 150 mM NaCl, 1% (v/v) Triton X-100, 5 mg/mL egg white lysozyme, and a protease inhibitor cocktail. After incubation at 37°C for 1 h, the solution was ultracentrifuged at 150,000 × *g* for 1h. If necessary, the pellets were resuspended with 10 mM HEPES-NaOH (pH7.5) and then ultracentrifuged again. The resultant pellets were resuspended in 100 μL of 10 mM HEPES-NaOH (pH7.5) and 150 mM NaCl.

For an ejection assay, the *S. lividans* strains harbouring pTYM19t::*sle1(HiBiT)-sle2* or pTYM19t::*sle1-sle2* were grown on solid Bennett’s-glucose media and cultivated at 30°C for 2 days. Mycelia were then scraped off the plate and treated as described above to extract SLPs. 15 μL of the isolated SLPs solution were mixed with the equal volume of 6 M urea or H_2_O and then 10 μL of this mixture was diluted with 90 μL of Nano-Glo luminescence reaction reagent (Promega Corporation, WI, USA). After equilibration to room temperature, luminescence was measured by a plate reader.

### Microscopy

For transmission electron microscopy, samples were attached to thin carbon film-coated TEM grids (ALLIANCE Biosystems, Osaka, Japan) and washed with H_2_O. The samples were then visualised by negative staining.

### Proteomic analysis

Approximately 10^4^ viable spores of each of the *S. lividans* strains (TK23, Δ*slpS*, and Δ*sle1/2*) were spread onto solid Bennett’s-glucose media and incubated at 30°C for 2 (mid-vegetative growth stage) or 3 (late-vegetative growth stage) days. The colonies were scraped off the plates and then disrupted by sonication in 50 mM HEPES-NaOH (pH7.5). The lysates were centrifuged at 10,000 × *g* for 3 min and the supernatants were used for further analyses. Three biologically independent samples were prepared for all the strains.

*E. coli* NiCo21(DE3) harbouring either of pET26b or pET26b::*3×FLAG-sle1-C* was precultured in 4 mL liquid LB-Lennox media containing 50 μg/mL kanamycin and then 200 μL of the cultures were inoculated into 100 mL of fresh liquid media containing kanamycin. After cultivation with shaking at 37°C for 3 h, 0.2 mM isopropyl-β-D-thiogalactopyranoside (IPTG) was added to each culture. After further cultivation at 30°C for 3 h, cells were harvested and disrupted by sonication in 10 mM HEPES-NaOH (pH7.5). The lysates were centrifuged at 5,000 × *g* for 3 min and the supernatants were used for further analyses. Three biologically independent samples were prepared for all the strains.

Proteins were separated by SDS–PAGE and were treated by in-gel digestion with trypsin. The digested samples were purified by Zip-tips, and were analysed by advance Nanoflor ultra-high performance liquid chromatography (Bruker, MA, USA) on a Q exactive quadrupole orbitrap mass spectrometer (Thermo Fisher) equipped with a Zaplous Column (0.2 i.d. × 50 mm; AMR, Inc., Japan, Tokyo) under the following conditions: column temperature, 35°C; mobile phase, gradient mixture of solvent A [0.1% formic acid] and solvent B [acetonitrile]; flow rate, 1.5 mL/min; and gradient elution, 0 min (solvent A:solvent B = 95:5), 20 min (35:65) and 21 mn (5:95). For protein identification, quantification, and comparison between two groups, database search was performed by label-free quantification workflow in Proteome Discoverer 2.5 inserted with a Sequest HT search engine with percolator against the genome of *S. lividans* and *E. coli*. Abundances of peptide spectral matches were averaged from the 2 technical replicates. Abundance ratios and q-values were calculated from the results for 3 biological replicates of each strain. The calculated abundance ratios of the detected proteins were automatically linked to the corresponding Gene Ontology terms for three categories (Biological Process, Cellular Component, and Molecular Function) and the proteins were grouped based on these values and terms.

### Protein expression and purification

*E. coli* NiCo21(DE3) cells harbouring the expression plasmids were precultured in liquid LB-Lennox medium containing 0.2-1% (w/v) glucose and 50 μg/mL ampicillin and then inoculated into 100 or 200 mL of fresh medium containing the antibiotics. After incubation at 37°C with shaking at 150 rpm, protein expression was induced by the addition of IPTG at the final concentration of 0.2 mM. For His_6_-SLIV_17110 (Sle2), the cultures were further incubated at 18°C for 18 h. For His_6_-maltose-binding protein (MBP)-Sle1, the cultures were further incubated at 30°C for 3 h. The cells were harvested by centrifugation and then disrupted by sonication in 5 mL of the lysis buffer comprising Tris-HCl buffer (pH8.0) and 10 mM imidazole. For purification, the filtered lysates were subjected to the Ni^2+^ affinity chromatography using His GraviTrap column. The eluates were concentrated and buffer-exchanged with 10 mM HEPES-NaOH (pH7.5) using Amicon Ultra-10K. Purified His_6_-MBP-Sle1 was further treated with TEV protease at 4°C for 20 h and then purified by the Ni^2+^ affinity chromatography as described above. The removal of the His_6_-MBP tag was confirmed by SDS–PAGE and purified tag-free Sle1 was concentrated and buffer-exchanged with 10 mM HEPES-NaOH (pH7.5) for further analysis.

### In vivo and in vitro protein expression assays

*In vivo* protein expression of the *E. coli* strains was analysed as follows. *E. coli* NiCo21(DE3) strains harbouring pBAD::*msfgfp* and either of pET26b or pET26b::*sle1-C* were precultured in 4 mL LB-Lennox media containing 50 μg/mL Kanamycin and ampicilin at 30°C overnight and the cultures with optical density (660 nm) 0.9 were inoculated to 100mL of fresh LB-Lennox media containing 50 μg/mL Kanamycin, 50 μg/mL ampicilin, and 0.05% (w/v) arabinose. These cultures were incubated in 96 well plates with shaking at 30°C for 80 min and then 0.1 mM IPTG was added. The cultures were further incubated with optical density at 660 nm and GFP fluorescence being measured every 20 min. To note, the detection settings for GFP fluorescence was adjusted to minimise the interference of background fluorescence through measuring the *E. coli* cultures without arabinose in prior to the above measurement.

*In vivo* protein expression of the *S. lividans* strains were analysed as follows. 10^6^ viable spores of the *S. lividans* strains harbouring a thiostrepton resistance gene and the P*_tipA_*-RS-*msfgfp* cassette were precultured in 4 mL Bennett’s-glucose media at 30°C for 2 days. After cultivation, 3 mL of a fresh medium was added to each of the cultures and 1 mL of the culture was poured into a 1.5 mL sterilised tube. After static incubation of the tubes at 30°C for 1 or 2 days, mycelia were concentrated by 10-fold by centrifugation at 3,000 × *g* for 5 min and the subsequent removal of 900 μL of the supernatant. 20 μL of the mycelial solution were added to 100 μL of a liquid Bennett’s-glucose medium in a 96 well plate with or without 20 μg/mL thiostrepton and 2 mM theophylline. After static preincubation at 30°C for 3 h, fluorescence was measured at the designated time points during incubation under the same condition. GFP fluorescence was calculated by subtracting the fluorescence of the culture without the inducers from that of the induced culture.

*In vitro* proteins expression was analysed as follows. The *E. coli* ribosome extract and other necessary components were derived from NEBExpress Cell-free *E. coli* Protein Synthesis System (New England Biolabs). The reaction mixture was comprised of 30S extract, T7 RNA polymerase, RNase A inhibitor, pET26b::*msfgfp*, and purified Sle1-C. This reaction mixture was separated into aliquots and incubated at 30°C. The reaction was stopped by cooling on ice at the designated time points. Protein expression was quantified by measuring GFP fluorescence of the samples.

### Protein interaction assays

*In vitro* protein interactions were assayed as follows.

The *E. coli* NiCo21(DE3) strains harbouring the pET26b::*3×FLAG-SLIV17115-N(sle1-N)*, pET26b::*3×FLAG-SLIV17115-C(sle1-C)*, pET26b::*FALG-slpT2*, pET26b::*FLAG-slp4*, and pET26b::*FLAG-slp5* were precultured in 4 mL of LB medium containing 50 μg/mL kanamycin at 30°C overnight. 200 μL of these precultures were inoculated into 4 mL of fresh LB medium containing 50 μg/mL kanamycin and incubated at 30°C for 1.5 h. 0.25 mM IPTG was added to the medium and further incubated at 30°C for 3 h. The *E. coli* NiCo21(DE3) strain harbouring pCold::*his6-SLIV_17110(sle2)* was precultured in 4 mL of LB medium containing 50 μg/mL ampicilin at 30°C overnight. 1 mL of the preculture was inoculated into 100 mL of LB medium containing 50 μg/mL ampicilin. After incubation at 30°C with shaking, the culture was cooled on ice and 0.2 mM IPTG was added. The culture was further incubated at 18°C overnight. The cultures were centrifuged and the collected cells were disrupted by sonication in 3 mL of 10 mM Tris-HCl and 10 mM imidazole (pH8.0). The lysates were cleared by filtration with 0.25 μm pore size. 700 μL (FLAG-SlpT2, FLAG-Slp4, and FLAG-Slp5) or 2 mL (3×FLAG-SLIV17115-N(sle1-N) and 3×FLAG-SLIV17115-C(sle1-C)) of the lysates containing the FLAG-tagged proteins were mixed with 2 mL of the lysate containing His_6_-SLIV_17110(Sle2). The mixture was incubated at room temperature for 25 min and then subjected to the Ni^2+^-affinity chromatography using His Gravitrap column. The inputs and eluates were analysed by western blot using the anti-DYKDDDDK antibody.

### Lipid membrane assays

Lipid membranes were purified as follows. *S. lividans* Δ*slpS* and *E. coli* NiCo21(DE3) harbouring pET26b::*3×FLAG-sle1-C* were grown on solid Bennett’s-glucose media and liquid LB-Lennox media at 30°C for 2 days and 3 h, respectively. The cells were collected and then disrupted by sonication in 10 mM HEPES-NaOH (pH7.5). After removing debris by a brief centrifugation, each of the supernatants were ultracentrifuged at 150,000 × *g* for 60 min. Each of the pellets containing the isolated lipid membranes were resuspended in 15% (w/v) iodixanol and then layered on top of an iodixanol gradient (20-50%) in a tube. After ultracentrifugation at 100,000 × *g* for 3 h, the iodixanol gradient was fractionated and lipid membrane content of each fraction was quantified by staining FM-143 dye and measuring its fluorescence. The lower density fractions containing lipid membranes were diluted with 10 mM HEPES-NaOH buffer (pH7.5) and then ultracentrifuged at 150,000 × *g* for 60 min. The resultant pellet was resuspended in 10 mM HEPES-NaOH (pH7.5) and used as purified lipid membranes for further analyses.

For analysing the interaction between lipid membranes and SLP, purified lipid membranes and SLPs were mixed and incubated at room temperature for 1h and then 4 °C overnight. To note, SLPs were extracted as described under *Extraction of SLPs* and washed with 10 mM HEPES-NaOH (pH7.5) to remove remaining detergent in the extract before use for the interaction assay with lipid membranes. The mixture was subjected to the iodixanol density-gradient ultracentrifugation as described above to separate lipid membranes and SLPs that are supposed to migrate to the lower and higher density fractions, respectively. The density gradient was fractionated and subjected to western blotting analysis, FM1-43 staining, and the luminescence assay.

### Bioinformatic analyses

Amino acid sequences of SLP-related proteins were retrieved from eCIStem (20) and the NCBI database. Protein structures were modeled using AlphaFold3 (51). Domain search and homology search were performed by the NCBI conserved domains database (52), the Basic Local Alignment Search Tool (BLAST) (53), and HHpred (54). Coiled-coil prediction was performed by WaggaWagga (55) and CoCoPRED (56). Transmembrane helix prediction was performed by TOPCONS (57).

### Western blotting

Rabbit antisera against SlpS, SlpT2, and Sle1 were developed using internal peptides RNDSERGVHKAPAN, CEGLSTQVEVEQRQEGGNNG, and SEANAATKRQRSSLEEAG, respectively, as antigens. Proteins were separated by SDS– PAGE with a 4-15% Mini-PROTEAN TGX Gel (Bio-Rad Laboratories, Inc., CA, USA) and then electroblotted onto a PVDF membrane. Transblotting was performed using Tris/glycine system with 10% (v/v) methanol or the Trans-Blot Turbo Transfer System (Bio-Rad Laboratories). After blocking with 5% (w/v) skim milk in Tris-buffered saline supplemented with 0.02% (v/v) Tween 20 (TBS-T buffer), the blots were incubated with each antibody (anti-SlpS, -SlpT2, and -Sle1 sera, diluted to 0.1%; anti-DYKDDDDK antibody [Proteintech, IL, USA], diluted to 0.01%) at room temperature for 60 min and horseradish peroxidase-conjugated secondary antibody (Goat anti-rabbit IgG H&L [HRP] ab6721[Abcam, Cambridge, UK], diluted to 0.01%) at room temperature for 45 min. If necessary, the primary antibodies were incubated with the blots at 4°C overnight. All the antibodies were diluted with the blocking buffer. The immunoreactive proteins were reacted with ImmunoStar LD (FUJIFILM Wako Chemicals, Osaka, Japan).

### Co-immunoprecipitation assay

*E. coli* NiCo21(DE3) harbouring either of pET26b::*sle1-C* or pET26b::*3×FLAG-sle1-C* was precultured in 4 mL liquid LB-Lennox media containing 50 μg/mL kanamycin and 0.5% (w/v) glucose and then these cultures were transferred to 100 mL LB-Lennox media containing kanamycin. These induction cultures were preincubated at 30°C for 30 min and then 0.3 mM IPTG was added to the cultures. After further cultivation for 5 h, the cells were harvested by a brief centrifugation and disrupted individually by sonication in a buffer comprising 10 mM HEPES-NaOH (pH7.4), 150 mM NaCl, and 0.5 mM EDTA. After 0.45 μm filtration, the protein extracts were mixed with DYKDDDDK Fab-Trap Agarose Beads (Proteintech) and interacting proteins were isolated following the protocol provided by the supplier. The proteins were eluted from the beads with Laemmli sample buffer containing 10% (v/v) 2-mercaptoethanol by incubating at 95°C for 5 min. The eluted proteins were separated by SDS–PAGE and CBB-visualised bands of interest were excised from the gel. The isolated proteins were digested with trypsin, and then the carbamidomethyl group was added to the -SH group of cysteine residues of the peptides. The resulting peptides were analysed by matrix-assisted laser desorption ionization-time of flight (MALDI-TOF) mass spectrometry. The obtained mass spectra derived from these peptides were analysed by a MASCOT database search (Matrix Science Ltd., London, UK). Proteins with the scores above the significance threshold were assigned to the bands.

### Growth measurements

The *E. coli* strains were precultured in 4 mL LB-Lennox media containing selection antibiotics and cultivated at 30°C until the cultures entered the stationary phase. They were inoculated into fresh media and incubated at 30°C in a 96 well plate with vigorous shaking and optical density at 660 nm and GFP fluorescence were measured by a plate reader during cultivation. For the functional assay of RS14790 and RS14785 in *E. coli*, the strains harbouring each of the plasmid sets derived from pET26b or pBAD/His A were precultured containing appropriate antibiotics and then the preculture was inoculated into fresh medium containing the antibiotics and inducers. The cultures were incubated at 30°C with shaking. Optical density was measured using a plate reader.

Growth of *S. lividans* was measured as follows. 10^6^ viable spores of the *S. lividans* strains were inoculated into 4 mL liquid Bennett’s-glucose media and cultivated at 30°C for 2 days. 1 mL of the preculture were inoculated into 100 mL of YPD medium. During cultivation with shaking (150 rpm) at 30°C, 1 mL of the cultures were sampled at the designated time points and their dry cell weights were measured immediately after sampling.

### Ribosome isolation

*S. lividans* TK23 was grown in 100 mL liquid Bennett’s-glucose medium and substrate mycelia were harvested by centrifugation. The mycelia were resuspended in 500 µL of cold resuspension buffer comprising 20 mM Tris-HCl (pH7.5), 15 mM MgCl_2_, and 1 mg/mL lysozyme. The resuspension solution was frozen and thawed three times to gently disrupt the cells. Next, 15 µL of 10% (w/v) sodium cholate and 10 µL of 1 mg/mL DNaseI were added, and the solution was centrifuged at 5,000 × *g* for 15 min to remove debris. The supernatant was placed on top of a sucrose-gradient solution in an ultracentrifugation tube. The sucrose-gradient solution was prepared by layering 0 and 50% (w/v) sucrose solutions containing 10 mM Tris-HCl (ph7.5), 50 mM NaCl, 50 mM KCl, 10 mM MgCl_2_, and 6 mM β-mercaptoethanol, and mixing the layered solutions using GRADIENT STATION (BioComp Instruments, Inc., New Brunswick, Canada). The sucrose-gradient solution was ultracentrifuged at 111,000 × *g* for 4 h and fractionated. UV adsorption was measured using a Triax Flow Cell (BioComp Instruments, Inc.) connected to the fractionator.

### Cell viability assay

10^6^ viable spores of the *S. lividans* strains were inoculated into 4 mL of Bennett’s-glucose media and cultivated at 30°C for 2 days. After cultivation, 3 mL of a fresh medium was added to each of the cultures and 300 μL of the culture and 200 μL of a fresh medium were mixed in a 1.5 mL sterilised tube. After static incubation of the tubes at 30°C for 1 or 2 days, 25 μL of the culture were mixed with 35 μL of a fresh medium and 40 μL of a luminescence reaction reagent reconstituted from RealTime-Glo MT Cell Viability Assay (Promega) in a 96 well plate. The reagent comprised 96 % (v/v) Bennett’s-glucose medium, 2% (v/v) NanoLuc solution, and 2% (v/v) substrate solution. The plate was incubated at 30°C without shaking and luminescence was measured every 20 min.

### Coculture assays

*S. lividans* strains harbouring a thiostrepton resistance gene were competed with other microorganisms with or without direct contact as follows. For the direct competition, approximately 3×10^4^ viable spores of *S. lividans* and competitor *Streptomyces* species were inoculated on solid MS media and cultivated at 30°C for 5 days. After cultivation, spores were collected from the cultures and then the numbers of viable spores of each strain were calculated by spreading the collected spores on solid TSB media with or without thiostrepton. For the indirect competition, approximately 10^4^ viable spores of the *S. lividans* strains were spotted onto a solid Bennett’s-glucose medium and the equivalent viable spores of competitor *Streptomyces* species or precultured *T. pulmonis* was spotted next to the *S. lividans* spores at a distance of 1 mm. The cultures were cultivated at 30°C.

## Data availability statement

The referred genome sequence of *S. lividans* TK24 is available at GenBank database (CP009124.1 [https://www.ncbi.nlm.nih.gov/nuccore/CP009124.1]). Any other datasets generated for the current study are available from the corresponding authors on request.

## Acknowledgements

T. Nagakubo was supported by a Grant-in-Aid for Scientific Research from the Japanese Society for the Promotion of Science (JSPS) (20J001208, 23K13863, and 25K21728) and Institute for Fermentation (Osaka, Japan). M. T. was supported by a Grant-in-Aid for Scientific Research from JSPS (23K26811) and the Suntory Rising Stars Encouragement Program in Life Sciences (SunRiSE). N. N. was supported by the Japan Science and Technology Agency (JPMJMI21G8 and JPMJGX23B2) and JSPS (23H05471). We thank Mitsue Arimoto and Dr. Koichiro Kako (University of Tsukuba) for their technical assistance and helpful suggestions for MALDI-TOF/MS analysis and ribosome isolation.

## Author contributions

T. Nagakubo conceived and designed the study. T. Nishiyama performed proteomic analysis. T. Nagakubo performed all other experiments. T. Nagakubo and T. Nishiyama analysed the data. T. Nagakubo, H. O, and M. T. drafted the manuscript. All the authors discussed the results and commented on the manuscript.

